# Multiplex logic processing isothermal diagnostic assays for an evolving virus

**DOI:** 10.1101/424440

**Authors:** Sanchita Bhadra, Miguel A. Saldaña, Hannah Grace Han, Grant L. Hughes, Andrew D. Ellington

## Abstract

We have developed a generalizable ‘smart molecular diagnostic’ capable of accurate point-of-care (POC) detection of variable nucleic acid targets. Our one-pot isothermal assay relies on multiplex execution of four loop-mediated isothermal amplification reactions, with primers that are degenerate and redundant, thereby increasing the breadth of targets while reducing the probability of amplification failure. An easy-to-read visual answer is computed directly by a multi-input Boolean OR gate signal transducer that uses degenerate strand exchange probes to assess any combination of amplicons. We demonstrate our platform by using the same assay to detect divergent Asian and African lineages of the evolving Zika virus (ZIKV), while maintaining selectivity against non-target viruses. Direct analysis of biological specimens proved possible, with 20 virions / µl being directly detected in human saliva within 90 minutes, and crudely macerated ZIKV-infected *Aedes aegypti* mosquitoes being identified with 100% specificity and sensitivity. The ease-of-use with minimal instrumentation, broad programmability, and built-in fail-safe reliability make our smart molecular diagnostic attractive for POC use.

---

While point-of-care (POC) diagnostic assays can be performed at or near the site of sample acquisition, they have for the most part been considered to be relatively simplistic tests that provide relatively little information to a clinician or public health worker. Familiar examples include electrochemical sensors for glucose,^1^ or rapid immunoassays for metabolites such as human chorionic gonadotropin (the canonical pregnancy test),^2^ and pathogens, either directly (influenza viruses) or via immune responses (antibodies against HIV-1/2).^3^ The range of conditions and pathogens that could be tested for could likely be greatly expanded by developing POC diagnostics for nucleic acids^3^ that would have greater sensitivity and accuracy. However, the current gold standard for molecular diagnostics, the quantitative polymerase chain reaction (qPCR), requires significant technical expertise and expensive and cumbersome equipment. Even portable instruments, such the Cepheid GeneXpert Omni, cost several thousand dollars and rely on expensive qPCR cartridge consumables for individual tests.

In contrast, isothermal nucleic acid amplification tests (iNAATs) have been developed that rival PCR in sensitivity, cost far less, and do not necessarily rely on complex instrumentation.^4^ To further the adoption of these assays, we and others have begun to develop ‘smart molecular diagnostics’ that can integrate information at the molecular level. For example, to increase signal specificity, a variety of sequence-specific probes, including molecular beacons, nuclease-dependent probes, and fluorescence resonance energy transfer (FRET) pairs have been adapted to isothermal amplification assays such as rolling circle amplification (RCA), recombinase polymerase amplification (RPA), or loop-mediated amplification (LAMP).^5, 6, 7^ More recently, RNA-guided CRISPR enzymes, such as Cas13a and Cas12a, have been used to signal the presence of isothermally generated amplicons.^8, 9^

In our own previous work, we developed oligonucleotide strand displacement (OSD) probes,^10^ that were triggered by strand exchange reactions with transiently single-stranded stem-loop sequences.^11^ This led to the exquisitely sensitive detection of LAMP amplicons without interference.^12^ One-pot LAMP-OSD assays that can work with crude samples are especially appealing for POC use.^13, 14^ These assays have been coupled to highly sensitive and reliable ‘yes/no’ output signals such as fluorescence, glucose, or hCG that can be readily read using off-the-shelf cellphones, glucometers, or pregnancy test strips, respectively.^12, 13, 15 16, 17, 18, 19^ Toehold switch RNA sensors have also been used to link iNAATs with *in vitro* reporter protein translation, leading to colorimetric signals.^20^

Even as these iNAAT tests move towards wider adoption, they all still face the same problem as other POC assays, in that the answers they give are relatively simplistic. However, given that the strand exchange reactions that underlie OSD probes were originally derived from far more complex DNA computations,^10, 21, 22, 23^ it may be possible to go beyond mere improvements in specificity and to integrate additional desirable features directly into the molecular diagnostic itself, such that the reaction helps to ‘compute’ its own outcome. As examples, we have previously developed strand exchange computation modules that can quantitate inputs to isothermal amplification reactions,^24^ or that can integrate multiple molecular signals via Boolean logic operations.^17^

We now attempt to take on real-world problems with computations that are embedded in the smart molecular diagnostics themselves. It is unfortunately a fundamental fact that diagnostic targets often evolve faster than assays designed to detect them. The resulting target sequence variations can easily prevent recognition of assay primers and probes, leading to test failure and false negative readouts, a problem which to date can only be solved via regular (and impractical) assay updates.^25, 26^ For instance, primers and probes for all eight published reverse transcription (RT) qPCR assays (qRT-PCR) for Zika virus (ZIKV) were found to have numerous mismatches with multiple ZIKV genomes sequenced from recent outbreaks.^27^ Some 20%-80% of ZIKV-infected patients are believed to have remained undiagnosed due to this incompatibility.^27, 28^

To overcome this problem and increase the overall accuracy of POC nucleic acid testing we have engineered additional computations into our smart molecular diagnostics that can compute the presence of almost any ZIKV variant. Strategic degenerate bases in 21 multiplexed primers allow the amplification of multiple ZIKV viral genes and gene variants in four, simultaneous LAMP reactions. The presence of any positive signal is calculated by multi-input Boolean ‘OR’-gated logic processors in a single one-pot reaction that directly analyzes crude biological samples and provides a visual yes/no readout for any ZIKV variant within 60-90 min. As with previous LAMP-OSD assays, the exquisite sequence-specificity of oligonucleotide strand exchange probes ensures signal specificity, while the built-in assay redundancy of the molecular logic processors guards against false negatives.

To demonstrate the utility of this smart molecular diagnostic we detect both Asian and African lineage ZIKV, which can differ by as much as 12%,^29^ and cleanly distinguish them from otherwise related and often co-circulating dengue virus (DENV) and chikungunya virus (CHIKV). As few as 100 ZIKV virions could be readily detected by direct analysis of artificially-infected human saliva. Pilot studies with un-infected and ZIKV-infected *Aedes aegypti* mosquitoes suggest that the integrated computations performed by our smart molecular diagnostic yield results that are on par with qRT-PCR, and for the first time demonstrate 100% specificity and sensitivity in a true POC format.

## Results

### Singleplex degenerate LAMP-OSD assays for detection of Asian and African lineage ZIKV

ZIKV has an ~10 kilobase RNA genome encoding a polyprotein that is cleaved into ten structural and non-structural (NS) proteins: 5’-Capsid (C)-preMembrane (prM)-Envelope (E)-NS1-NS2A-NS2B-NS3-NS4A-NS4B-NS5-3’. ZIKV clusters into two African lineages and one Asian lineage that includes the current epidemic strains.^29^ The Asian and African lineages of ZIKV show 0.2% to 10.6% intra-lineage and 4.5% to 12.1% inter-lineage nucleotide variation.^29^ The epidemic strains have accumulated additional changes and have potentially undergone multiple natural recombinations.^29, 30^ The strains are likely continuing to evolve, presenting a challenging target for surveillance and diagnostics. Here, we designed four reverse transcription (RT) LAMP assays to amplify relatively conserved regions in four ZIKV genes – capsid (CA), NS1, NS3, and NS5. Each individual assay included two inner primers (FIP and BIP), two outer primers (B3 and F3), and one loop primer (LP). The CA LAMP assay utilized an additional sixth primer, termed stem primer^31^ (SP), complementary to the target region between F1 and B1 priming sites (**Figure 1A**). Specific nucleotide positions within these primers were substituted with degenerate nucleobases to allow pairing with variant ZIKV RNA (**Supplementary Table T1**). The NS3 LAMP primer set had the lowest degeneracy with only 10 positions varying between two nucleobases and one position varying between three nucleobases. Twelve positions in the NS1 LAMP primer set varied between two nucleobases. The CA LAMP primers had a slightly higher degeneracy with 13 polymorphic positions varying between two bases. The NS5 primer set was the most degenerate with 15, 6, and 1 positions varying between two, three, or four bases, respectively.

**Figure 1.**
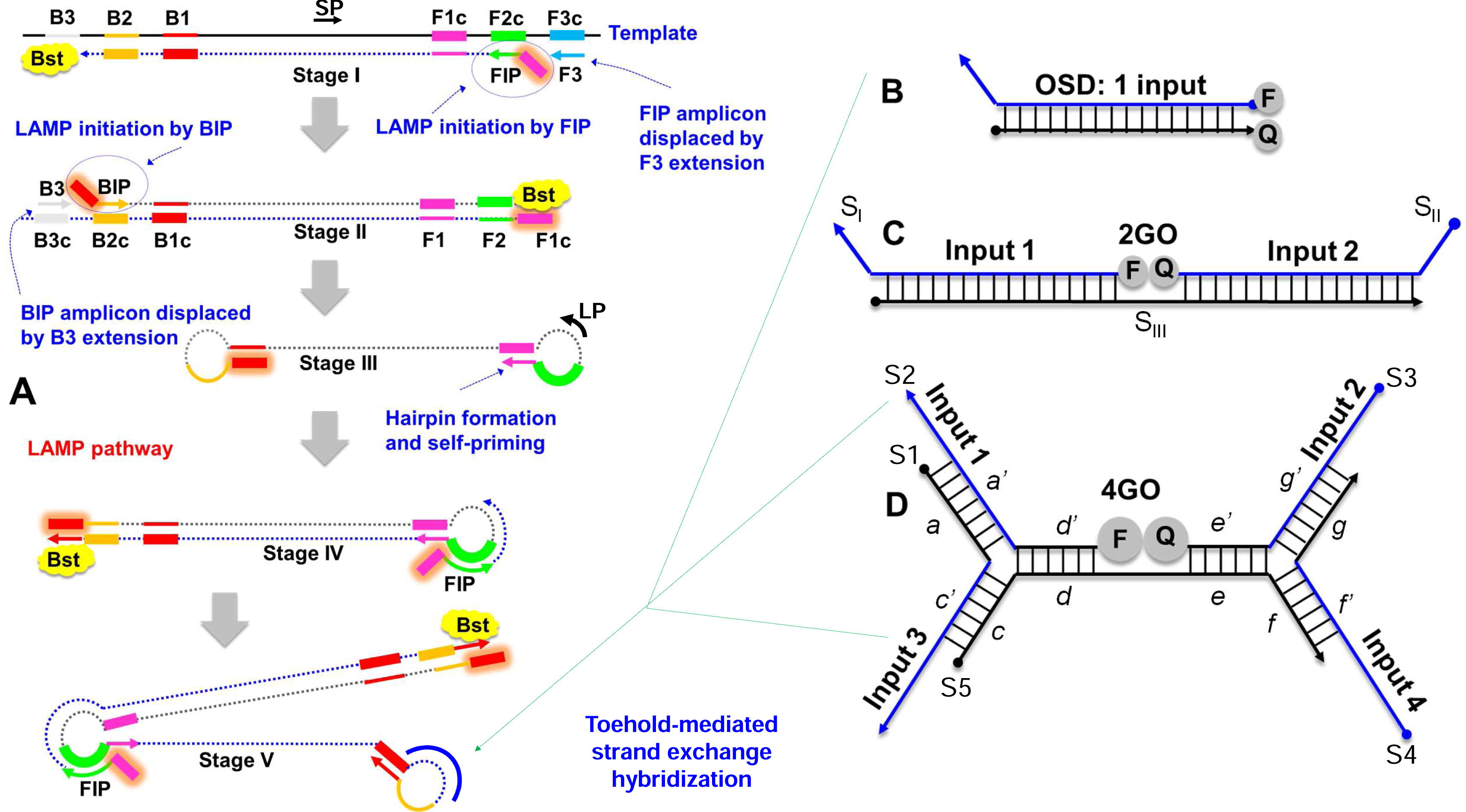
Schematic depicting (A) loop-mediated isothermal amplification (LAMP) integrated with (B) one-, (C) two-, or (D) four-input oligonucleotide strand exchange signal transducers. LAMP uses 2 inner (FIP and BIP) and 2 outer (F3 and B3) primers along with the optional stem (SP) and loop (LP) primers to prime strand displacement DNA amplification by Bst DNA polymerase. The resulting continuous amplification (initiated by both new primer-binding and by self-priming) generates double-stranded concatameric amplicons containing single-stranded loops to which non-priming oligonucleotide strand exchange signal transducers can hybridize. The one-input OSD signal transducer composed of one long and one short DNA strand can hybridize to a single LAMP amplicon loop sequence leading to separation of the fluorophore (F) and quencher (Q). The OR Boolean logic processing two-input strand exchange transducer, 2GO, is composed of two labeled strands, S_I_ and S_II_, and a third bridging strand S_III_. Either S_I_ and/or S_II_ can hybridize to their specific LAMP loop sequences resulting in separation of F and Q. The four-input 4GO probe composed of 5 DNA strands (S1-S5) can hybridize to any combination of up to four different LAMP amplicon loops and perform an OR Boolean operation to produce fluorescence signal. 4GO probe is denoted in terms of lettered domains (*a-g*), each of which represents a short fragment of DNA sequence in an otherwise continuous oligonucleotide strand. Complementarity is denoted by a single prime symbol.

To ensure readout specificity, individual hemiduplex OSD probes^12^ were designed for each of the four ZIKV LAMP amplicons (**Figure 1B**). To facilitate detection of LAMP amplicons from different viral lineages, probe nucleotide positions corresponding to polymorphic target loci were substituted with degenerate bases (**Supplementary Table T2**). The NS3 OSD probe was the least degenerate with only two positions in the long and the short probe strands varying between two bases. The capsid OSD probe contained two nucleotide variations at four and two positions in the long and the short strand, respectively. Three positions in both the long and the short strands of the NS5 OSD probe varied between two bases. The NS1 OSD probes displayed the greatest degeneracy with four positions in both the long and the short strands varying between two bases.

Performance of these four degenerate reverse transcription LAMP-OSD assays was optimized using singleplex reactions containing different amounts of synthetic target RNA derived from Asian lineage ZIKV (**Supplementary Table T3**). All four assays could detect a few hundred copies of synthetic target RNA within 60 min without producing spurious signal (**Figure 2**). Furthermore, the high signal amplitude and low noise allowed simple yes/no visual readout and cellphone imaging of these assays – target RNA yielded bright visible fluorescence while assays lacking specific templates remained dark (**Figure 2**).

**Figure 2.**
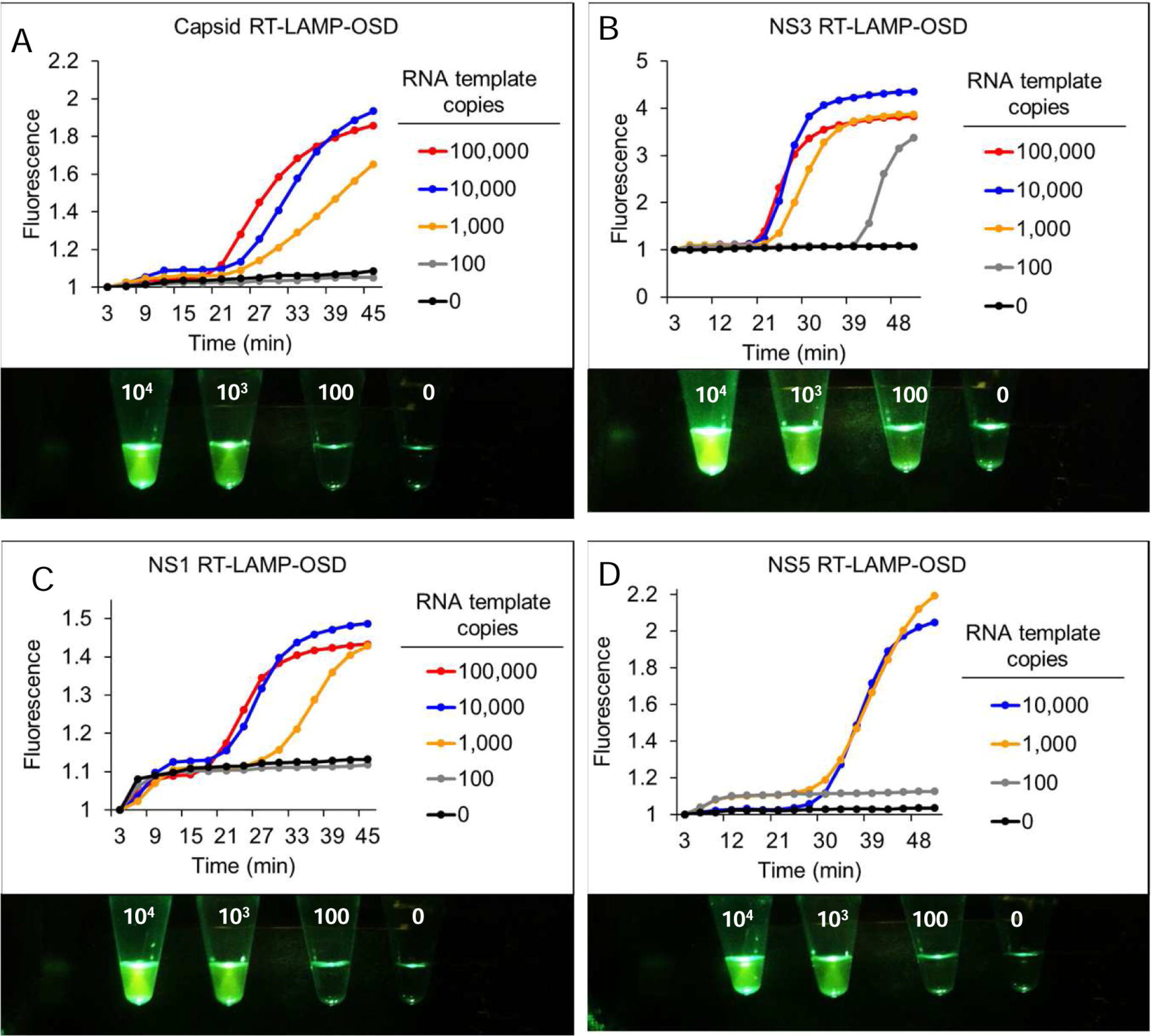
Detection of Zika virus capsid, NS1, NS3, and NS5 genes using real-time and visually-read reverse transcription LAMP-OSD assays. Indicated copies of capsid (A), NS3 (B), NS1 (C), and NS5 (D) synthetic RNA templates were amplified by singleplex degenerate LAMP-OSD assays specific to each template. OSD fluorescence signals measured in real-time during LAMP amplification are depicted as red (10^5^ template copies), blue (10^4x^ template copies), orange (10^3^ template copies), gray (100 template copies), and black (0 template copies; 10^6^ non-template RNA) traces. The x-axis depicts the duration of LAMP amplification. OSD fluorescence was also imaged at amplification endpoint using a cellphone (images depicted at the bottom of each panel). Numbers on each assay tube in these images indicate the RNA template copies used. Representative results from three replicate experiments are depicted.

To verify that all the four assays could identify both Asian and African lineage ZIKV genomes we individually challenged singleplex NS1, NS3, NS5, and CA LAMP-OSD assays with genomic RNA from 9 Asian ZIKV, and 2 African ZIKV strains (**Supplementary Table T4**). Reaction specificity was assessed by seeding duplicate assays with genomic RNA of related dengue virus (DENV) serotypes 1-4 and chikungunya virus (CHIKV) that often co-circulate with ZIKV.^32^ Endpoint OSD fluorescence in all LAMP assays seeded with Asian or African ZIKV genomes was significantly elevated above background noise in assays seeded with DENV or CHIKV genomic RNA (**Figure 3**). These results demonstrate that all four degenerate LAMP-OSD assays were capable of lineage-independent ZIKV detection without cross-reaction with or interference from non-target nucleic acids.

**Figure 3.**
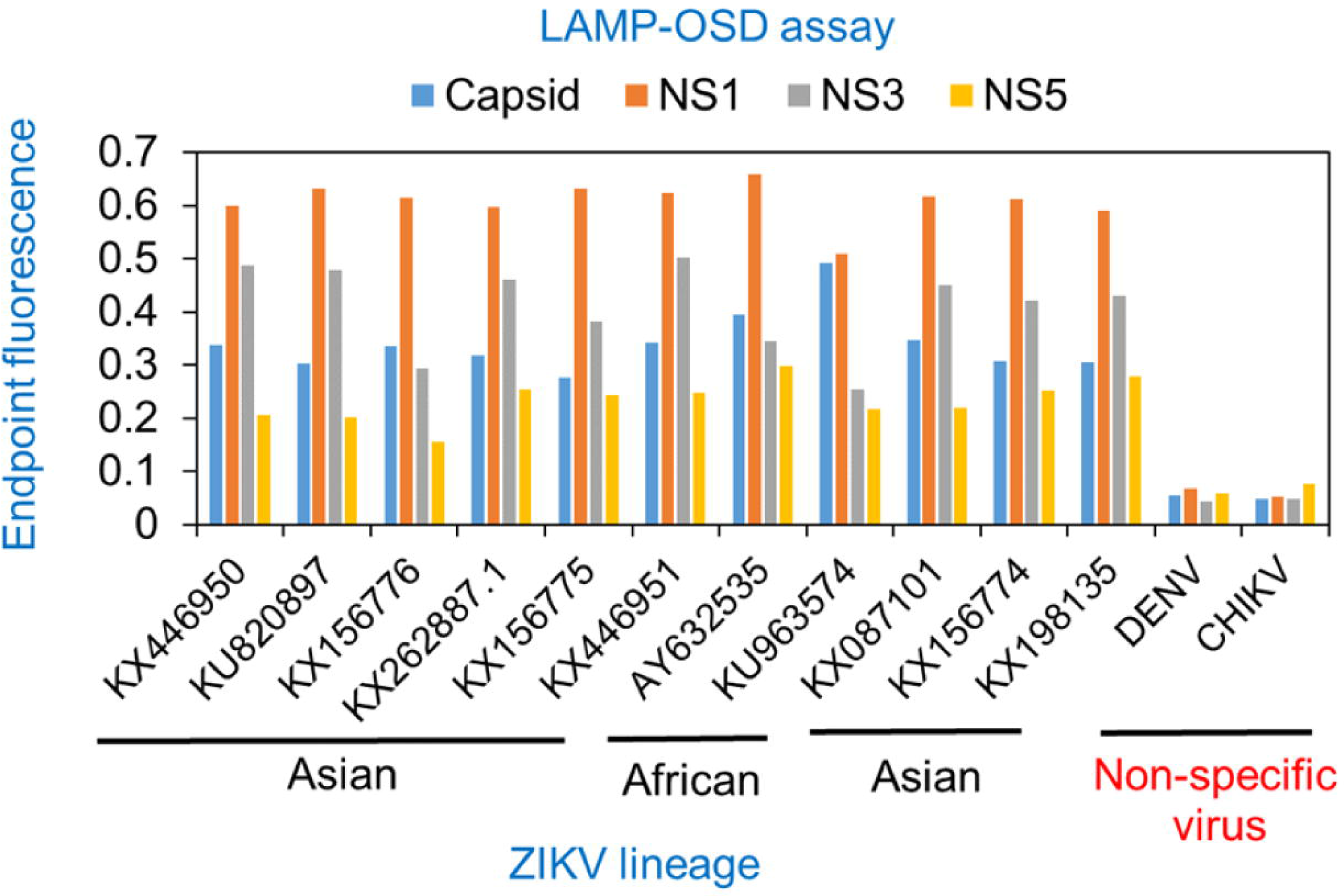
Detection of Asian and African lineage ZIKV using singleplex degenerate reverse transcription LAMP-OSD assays. Whole cell RNA purified from cells infected with DENV, CHIKV, or Asian or African lineage ZIKV (indicated by their GenBank accession numbers) were used as templates for amplification in singleplex degenerate RT-LAMP-OSD assays for Zika virus capsid, NS1, NS3, and NS5 genes. OSD fluorescence signals measured at amplification endpoint using LightCycler 96 real-time PCR machine are depicted as blue (capsid), orange (NS1), gray (NS3), and yellow (NS5) bars. Representative results from three replicate experiments are depicted.

### Multiplexed LAMP with degenerate primers and OSD probes

To create a multiplexed, internally redundant assay that simultaneously amplifies and detects CA, NS1, NS3, and NS5 sequences, all 21 LAMP primers and 4 hemi-duplex OSD reporters were combined in a single reaction (4-Plex-LAMP-OSD). To verify that each individual reverse transcription LAMP-OSD reaction was functional in this multiplex environment, replicate multiplex assays were supplemented with all 21 primers but only a single type of OSD reporter at a time. In the presence of all four ZIKV synthetic RNA templates, multiplexed 4-Plex-LAMP-OSD demonstrated exponential fluorescence accumulation. Despite co-mingling of multiple degenerate primers and OSD probes, no spurious signals were observed in the absence of specific templates (**Figure 4A**). As a result, brightly fluorescent ZIKV-positive assays containing only a few hundred copies of ZIKV templates could be readily distinguished from dark ZIKV-negative reactions by simple visual examination (**Figure 4B**). Each individual LAMP assay was functional in the multiplex assay, as indicated by accumulation of fluorescence in 4-Plex-LAMP-OSD reactions probed with individual OSD reporters (**Figures 4C, D, E, F**). The multiplexed 4-Plex-LAMP-OSD assay could also readily identify the more complex ZIKV genomic RNA from both Asian and African lineages without cross-reaction with DENV and CHIKV genomic nucleic acids (**Figure 4G**).

**Figure 4.**
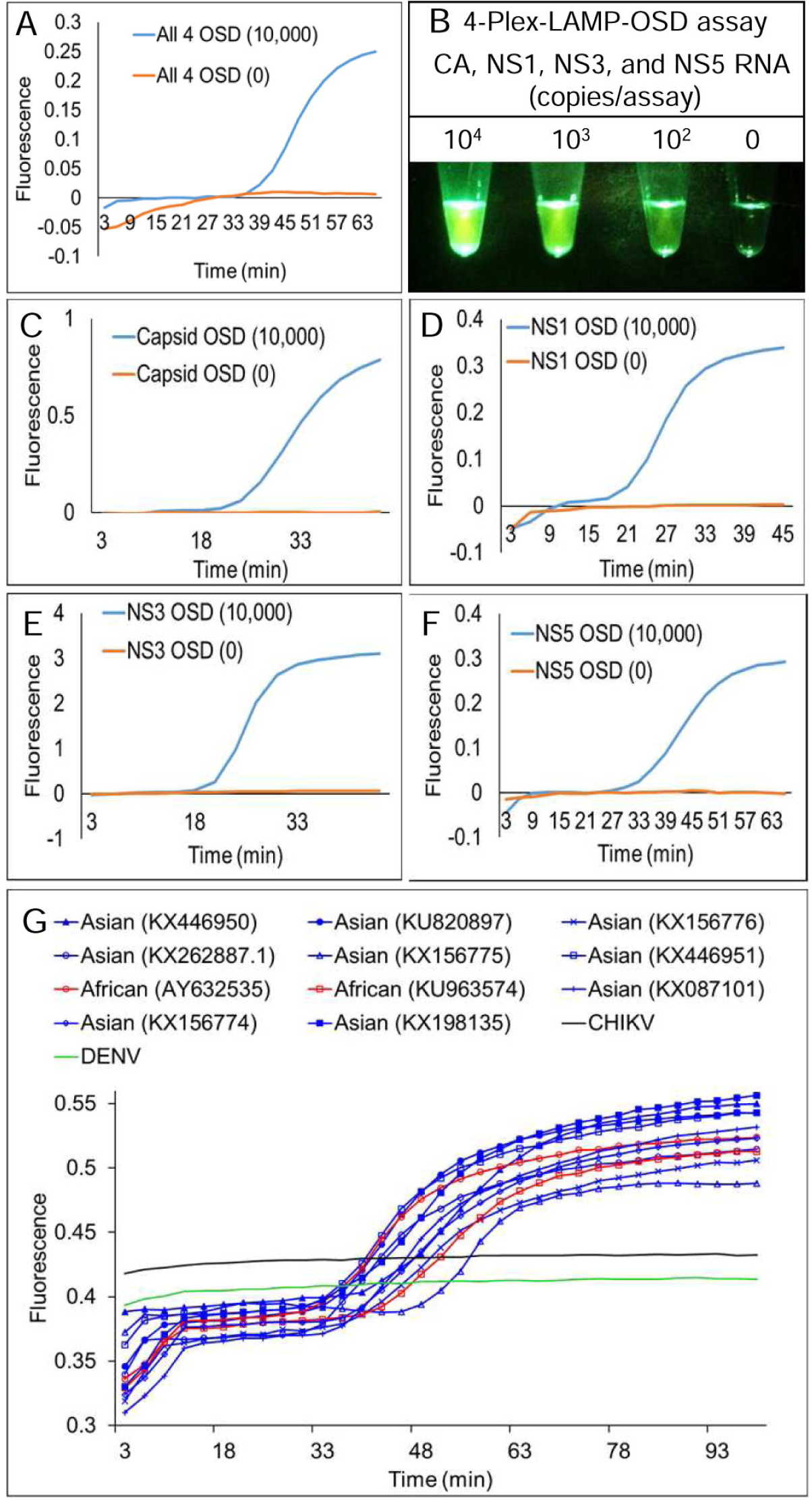
Simultaneous detection of four Zika virus genes using multiplex reverse transcription degenerate LAMP-OSD (4-Plex-LAMP-OSD) assay. (A) Real-time 4-Plex-LAMP-OSD – synthetic RNA mixtures containing indicated copies of each of the four ZIKV synthetic RNA templates (CA, NS1, NS3, and NS5) were amplified using 4-Plex-LAMP-OSD assays containing 21 degenerate primers and 4 degenerate OSD probes for simultaneous LAMP amplification and sequence-specific detection of all four ZIKV targets. OSD fluorescence signals measured in real-time during LAMP amplification are depicted as blue (10,000 copies each of CA, NS1, NS3, and NS5 RNA) and orange (0 ZIKV RNA; 10^6^ copies of DENV RNA) traces. The x-axis depicts the duration of LAMP amplification. (B) Endpoint 4-Plex-LAMP-OSD assay with visual detection – synthetic RNA mixtures containing indicated copies of each of the four ZIKV synthetic RNA templates (CA, NS1, NS3, and NS5) were amplified using degenerate 4-Plex-LAMP-OSD assays. OSD fluorescence was imaged after 90 min of amplification using a cellphone. Numbers above each assay tube indicate the RNA template copies used. The reaction with ‘0’ ZIKV RNA received 10^6^ copies of DENV RNA. (C-F) Performance of individual assays in the 4-Plex-LAMP-OSD system – synthetic RNA mixtures containing indicated copies of each of the four ZIKV synthetic RNA templates (CA, NS1, NS3, and NS5) were amplified using 4-Plex-LAMP-OSD assays containing LAMP primers for all four targets but only one type of OSD for either capsid (C), NS1 (D), NS3 (E), or NS5 (F) amplicons. OSD fluorescence signals measured in real-time during LAMP amplification are depicted as blue (10,000 copies each of CA, NS1, NS3, and NS5 RNA) and orange (0 ZIKV RNA; 10^6^ copies of DENV RNA) traces. The x-axis depicts the duration of LAMP amplification. (G) Detection of Asian and African lineage ZIKV genomic RNA using degenerate 4-Plex-LAMP-OSD assays. Whole cell RNA from cells infected with DENV, CHIKV, or Asian or African lineage ZIKV (indicated by their GenBank accession numbers) were used as templates for amplification. Real-time OSD fluorescence signals are depicted as blue (Asian), red (African), black (CHIKV), and green (DENV) traces. The x-axis depicts the duration of LAMP amplification. For all experiments, representative results from three replicate tests are depicted.

### Multiplexed LAMP with degenerate primers and four-input logic-processing probes

After functionally verifying the feasibility of multiplexed reverse transcription LAMP-OSD using degenerate primers and OSD probes we sought to reduce reaction cost and enhance assay versatility by replacing the four individual OSD probes with a single four-input OR Boolean logic-processing strand exchange probe (4GO) that would ‘compute’ the presence of ZIKV by simultaneously looking across CA, NS1, NS3, and NS5 amplicons for similarity to the probe. Unlike four individual OSDs that required a combination of 8 fluorophore or quencher-labeled oligonucleotides, the three-way junction 4GO probe would need a single fluorophore-quencher pair (**Figure 1**). If either CA, NS1, NS3, or NS5 amplicon was similar enough to initiate strand exchange, the fluorophore would separate from the quencher and the 4GO probe would light up.

The 4GO probe was composed of 5 oligonucleotides (S1 – S5) with a total of 19 degenerate loci that varied between two nucleobases (**Supplementary Table T2**). Oligonucleotides S1 – S4, designed based on validated degenerate OSD sequences, served as strand exchange probes for NS1, NS5, CA, and NS3 amplicons, respectively. The S5 strand acted as a scaffold for 4GO probe assembly (**Figure 1D**). Domain *c* at the 5’-end of S5 was complementary to domain *c’* in S1, while, domain *f* at the 3’-end of S5 was complementary to domain *f’* of the S4 strand. The intervening region of S5 contained two short target-independent sequences – domain *d* complementary to domain *d’* at the fluorophore-labeled 5’-end of S2 and domain *e* complementary to domain *e’* at the quencher-labeled 3’-end of S3. Concomitant hybridization of S2 and S3 to S5 would juxtapose the fluorophore and quencher leading to loss of signal. However, at temperatures ≥ 20 °C, pairing of S2 and S5 was contingent upon formation of a three-way junction with S1 via interactions between complementary S2 domain *a’* with S1 domain *a* and S5 domain *c* with S1 domain *c’*. Similarly, stable interaction of S3 and S5 was dependent upon a three-way hybridization with S4. As a result, strand exchange between the 4GO probe and any one or more of the ZIKV amplicons would separate the fluorophore-bearing S2 from quencher-labeled S3. For example, the NS5 LAMP amplicon loop would directly remove S2 from the 4GO probe. Meanwhile, by binding S1, the NS1 amplicons would destroy the three-way junction between S1, S2, and S5. Consequently, the 4GO probe would transduce the presence of one or any combination of the four ZIKV amplicons into a single-channel endpoint fluorescence that could be easily read at ambient temperature.

To test 4GO probe function, multiplex reverse transcription LAMP reactions containing all 21 degenerate primers and the degenerate 4GO probe (4-Plex-LAMP-4GO) were spiked with a single type or a combination of all four of the ZIKV synthetic RNA templates NS1, NS3, NS5, and CA. As a control to assess signal specificity, duplicate reactions were assembled using a non-specific RNA template and its cognate LAMP primers that would lead to generation of non-specific LAMP amplicons. Following LAMP amplification, 4GO probe fluorescence was found to be elevated in all reactions containing even a few hundred copies of at least one of the ZIKV templates (**Figures 5A-E**). This ZIKV-specific signal remained consistently above noise generated in the absence of specific amplicons. These results demonstrate that the 4GO probe was able to function specifically in multiplex LAMP to identify and signal the presence of a few hundred copies of one or all four ZIKV amplicons without interference from non-specific templates or amplicons.

**Figure 5.**
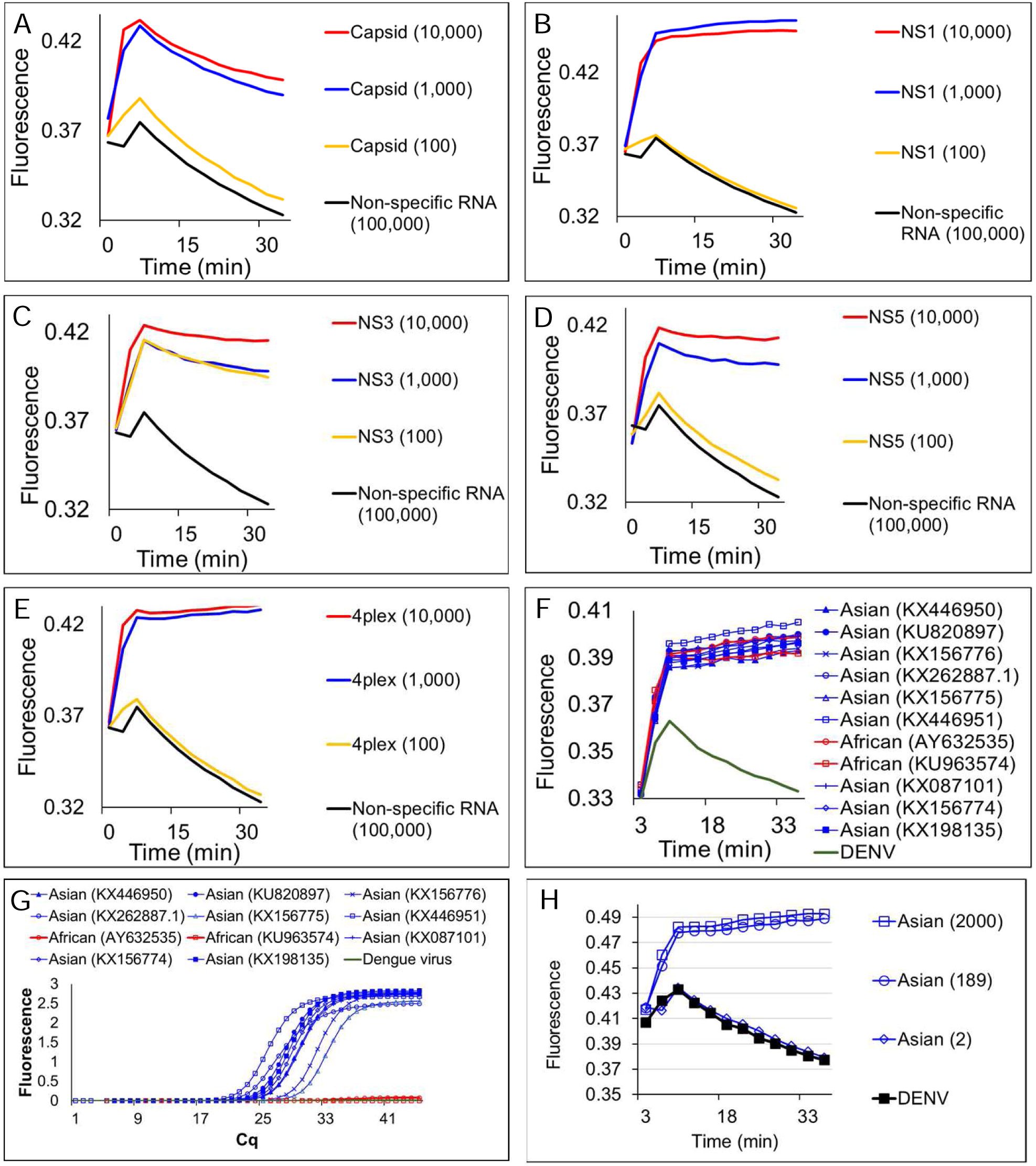
Simultaneous detection of four Zika virus genes using degenerate 4GO probes and multiplex degenerate reverse transcription LAMP (4-Plex-LAMP-4GO) in one-pot assays. Indicated copies of capsid, NS1, NS3, and NS5 synthetic target RNA were amplified either individually (panels A, B, C, and D, respectively) or as a mixture (panel E) using 4-Plex-LAMP-4GO assays containing LAMP primers for all four ZIKV targets and the four-input 4GO probe. 4GO probe fluorescence, measured in real-time at 37 °C after 90 min of LAMP amplification, is depicted as red (10,000 template copies), blue (1,000 template copies), yellow (100 template copies), and black (non-specific LAMP primers with 10^5^ copies of its target RNA) traces. The x-axis depicts the duration of endpoint signal measurement. (F) Detection of Asian and African lineage ZIKV genomic RNA using degenerate 4-Plex-LAMP-4GO assays. Whole cell RNA from cells infected with DENV, or Asian or African lineage ZIKV (indicated by their GenBank accession numbers) were used as templates for amplification. 4GO probe fluorescence, measured in real-time at 37 °C after 90 min of LAMP amplification, is depicted as blue (Asian), red (African), and green (DENV) traces. The x-axis depicts the duration of endpoint signal measurement. (G) Detection of Asian and African lineage ZIKV genomic RNA using TaqMan qRT-PCR assay specific for Asian lineage ZIKV NS2b gene. Same amount of viral genomic RNA as was used in panel F were amplified and real-time measurements of assay fluorescence are depicted as blue (Asian), red (African), and green (DENV) traces. (H) Detection limit of degenerate 4-Plex-LAMP-4Go assay for ZIKV genomic RNA. Indicated copies of an Asian lineage ZIKV genome or non-specific DENV genomes were amplified using 4-Plex-LAMP-4GO assays. 4GO probe fluorescence, measured in real-time at 37 °C after 90 min of LAMP amplification, is depicted as blue (Asian) and black (DENV) traces with template copies indicated by open squares (2000 genomes), open circles (189 genomes), and open diamonds (2 genomes). The x-axis depicts the duration of endpoint signal measurement. For all experiments, representative results from three replicate tests are depicted.

4GO probes were also able to specifically identify all 9 Asian and both African ZIKV strains tested without cross reaction with related viruses (**Figure 5F**). Endpoint fluorescence signals in all degenerate 4-Plex-LAMP-4GO assays seeded with Asian or African lineage ZIKV genomic RNA were consistently elevated above background noise in assays containing non-specific DENV genomes. In contrast, a previously reported Asian ZIKV NS2b gene-specific TaqMan qRT-PCR assay^33^ was able to detect only Asian lineage viral genomes while failing to amplify African lineage genomic RNA (**Figure 5G**). Asian genomic RNA copy numbers estimated from this qRT-PCR assay (**Supplementary Figure 1**) suggested that 4-Plex-LAMP-4GO assays could readily detect a few hundred copies of viral genomes (**Figure 5H**). These results demonstrate that a single one-pot assay comprised of 21 degenerate LAMP primers and a five-stranded degenerate logic processing strand exchange probe could reliably detect viral variants within a genus and distinguish them from related viruses. Furthermore, by integrating signals from four separate inputs into one output via a single fluorophore-quencher pair, the five-stranded 4GO probe could capture the viral diversity at a fraction of the cost of four individually-labeled OSD probes.

### Multiplexed LAMP with degenerate primers and two-input logic-processing probes

Our results demonstrated that the 4-Plex-LAMP-4GO assay could specifically identify both Asian and African ZIKV without non-specific signaling. However, a fluorimeter was necessary for assay readout due to the relatively high background noise of the 4GO probe. Therefore, we sought to further engineer the logic processing probe in order to achieve a signal-to-noise ratio that could be visually discriminated and thereby allowed easy ‘yes/no’ assay readout in austere conditions without complex instruments. We surmised that the fluorophore was insufficiently quenched in the five-stranded 4GO probe due to its high degeneracy level. To reduce probe degeneracy, we engineered a bipartite four-input signal processor composed of two OR gated strand exchange modules (2GO) (**Figure 1C**). Each 2GO probe was comprised of 3 degenerate oligonucleotides (S_I_-S_III_) that were partially complementary to each other. The 5’-ends of both S_I_ strands were labeled with the same type of fluorophore while the 3’-ends of S_II_ strands were labeled with the corresponding quencher molecules. Simultaneous hybridization of S_I_ and S_II_ to the unlabeled S_III_ strand would juxtapose the fluorophore and quencher resulting in loss of signal. Short single-stranded toeholds at the 3’- and 5’-ends of S_I_ and S_II_, respectively could independently initiate strand exchange with their cognate target sequences resulting in separation of the fluorophore from the quencher and a concomitant rise in signal. For instance, the CAN3.2GO probe fluorescence would increase if it underwent strand exchange with either CA or N3 amplicons. Similarly, the N1N5.2GO probe would signal if either or both NS1 and NS5 amplicons were present.

To functionally verify the bipartite signal processor, reverse transcription LAMP assays containing the degenerate CAN3.2GO or the NS1NS5.2GO probes were supplemented with primers specific to either ZIKV NS1, NS3, NS5, or CA genes. At amplification endpoint, fluorescence of CAN3.2GO probe increased only in assays containing CA or N3 amplicons (**Figure 6A**). Meanwhile the N1N5.2GO probe was activated only in the presence of NS1 or NS5 sequences. Both probes remained quenched in the presence of non-specific LAMP amplicons. Moreover, amplicon-specific 2GO probe fluorescence could be readily distinguished from non-specific noise simply by direct visual examination of assay tubes or their cellphone images (**Figure 6B**).

**Figure 6.**
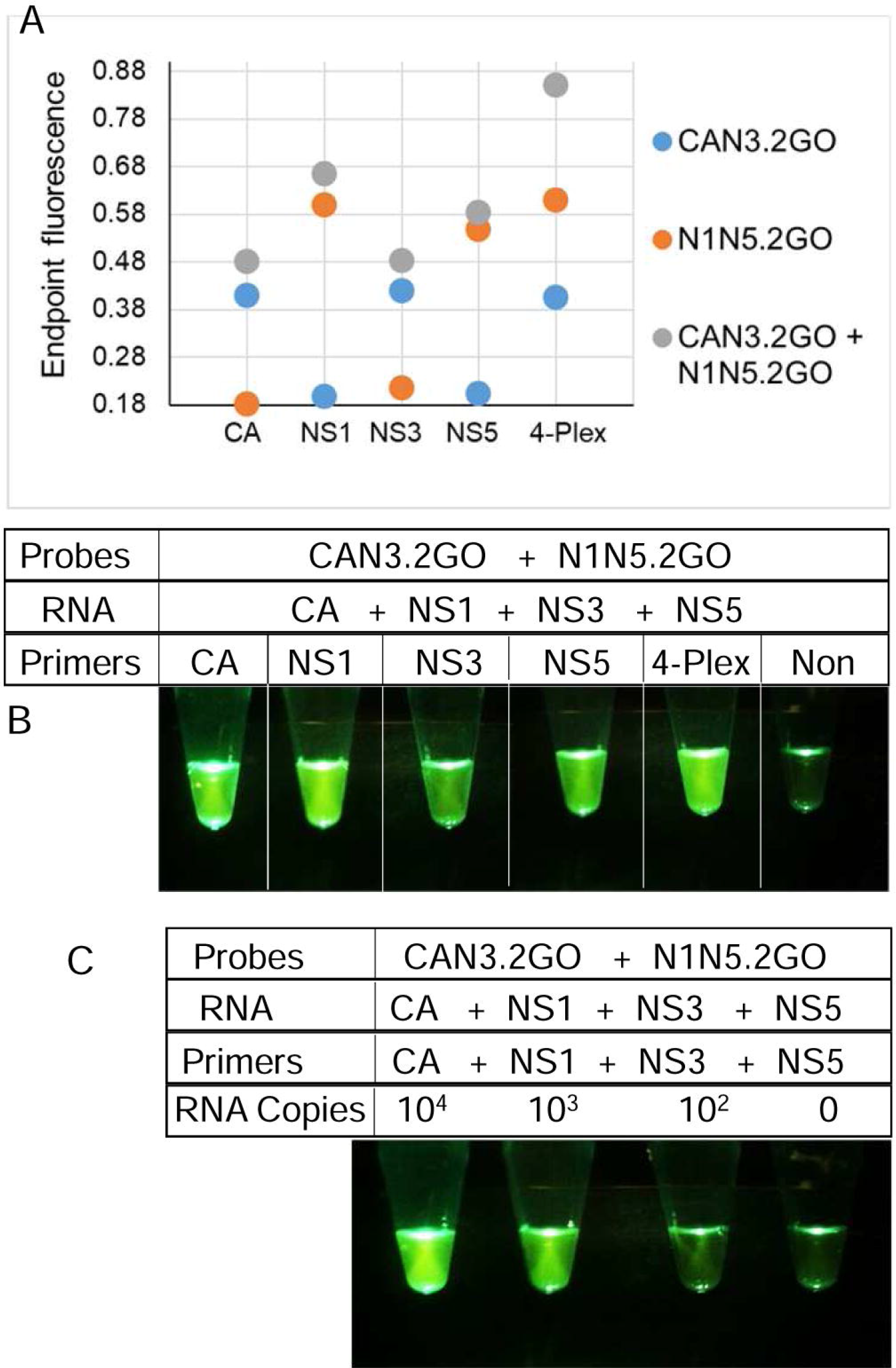
Detection of ZIKV RNA using two-input 2GO probes and degenerate reverse transcription LAMP. (A) Sequence-dependent activation of 2GO probes – synthetic RNA mixtures of 10^6^ copies of CA, NS1, NS3, and NS5 RNA were amplified using single- or 4-plex degenerate LAMP assays containing either one or both CAN3.2GO and N1N5.2GO probes. 2GO probe fluorescence signals measured at amplification endpoint using LightCycler 96 real-time PCR machine are depicted as blue (LAMP with only CAN3.2GO), orange (LAMP with only N1N5.2GO), and gray (LAMP with both CAN3.2GO and N1N5.2GO) dots. LAMP primer specificities are indicated on the x-axis. (B) Visual readout of degenerate LAMP-2GO assays. Cellphone image depicts 2GO probe fluorescence at amplification endpoint in single- or 4-plex ZIKV LAMP assays containing both CAN3.2GO and N1N5.2GO probes and 10^6^ copies of all four synthetic ZIKV RNA and a non-specific LAMP assay (‘Non’) containing its cognate RNA. (C) Detection limit of visually-read degenerate 4-Plex-LAMP-2GO assays. Cellphone image depicts endpoint 2GO probe fluorescence of multiplex degenerate RT-LAMP assays containing primers and indicated template RNA copies of all four ZIKV targets. The reaction without any ZIKV RNA contained a non-specific RNA and its cognate LAMP primers. For all experiments, representative results from three replicate tests are depicted.

The complete multiplex assay system (4-Plex-LAMP-2GO) containing 21 degenerate LAMP primers and the bipartite signal processor (CAN3.2GO + N1N5.2GO) was tested using different amounts of a synthetic RNA mixture containing all four ZIKV templates (NS1, NS3, NS5, and CA). Assays containing only a few hundred copies of ZIKV RNA produced bright fluorescence that could be readily captured using unmodified smartphone camera (**Figure 6C**). In contrast, assays containing non-specific amplicons remained dark thereby allowing easy visual distinction from ZIKV-positive assays.

Similar to LAMP-OSD and 4-Plex-LAMP-4GO assays, 4-Plex-LAMP-2GO assays could also detect all nine Asian and both African ZIKV strains tested while suppressing spurious signal from dengue viruses (serotypes 1-4) (**Figure 7A**). Examination of endpoint smartphone images of 4-Plex-LAMP-2GO assays revealed that as few as 189 copies of a representative Asian ZIKV genomic RNA (quantified using NS2b-specific TaqMan qRT-PCR) could be reliably identified using simple yes/no visual readout of presence or absence of fluorescence (**Figures 7B and Supplementary Figure 1**). Similarly, African ZIKV genomic RNA could also be detected using smartphone imaged 4-Plex-LAMP-2GO assays (**Figure 7B**).

**Figure 7.**
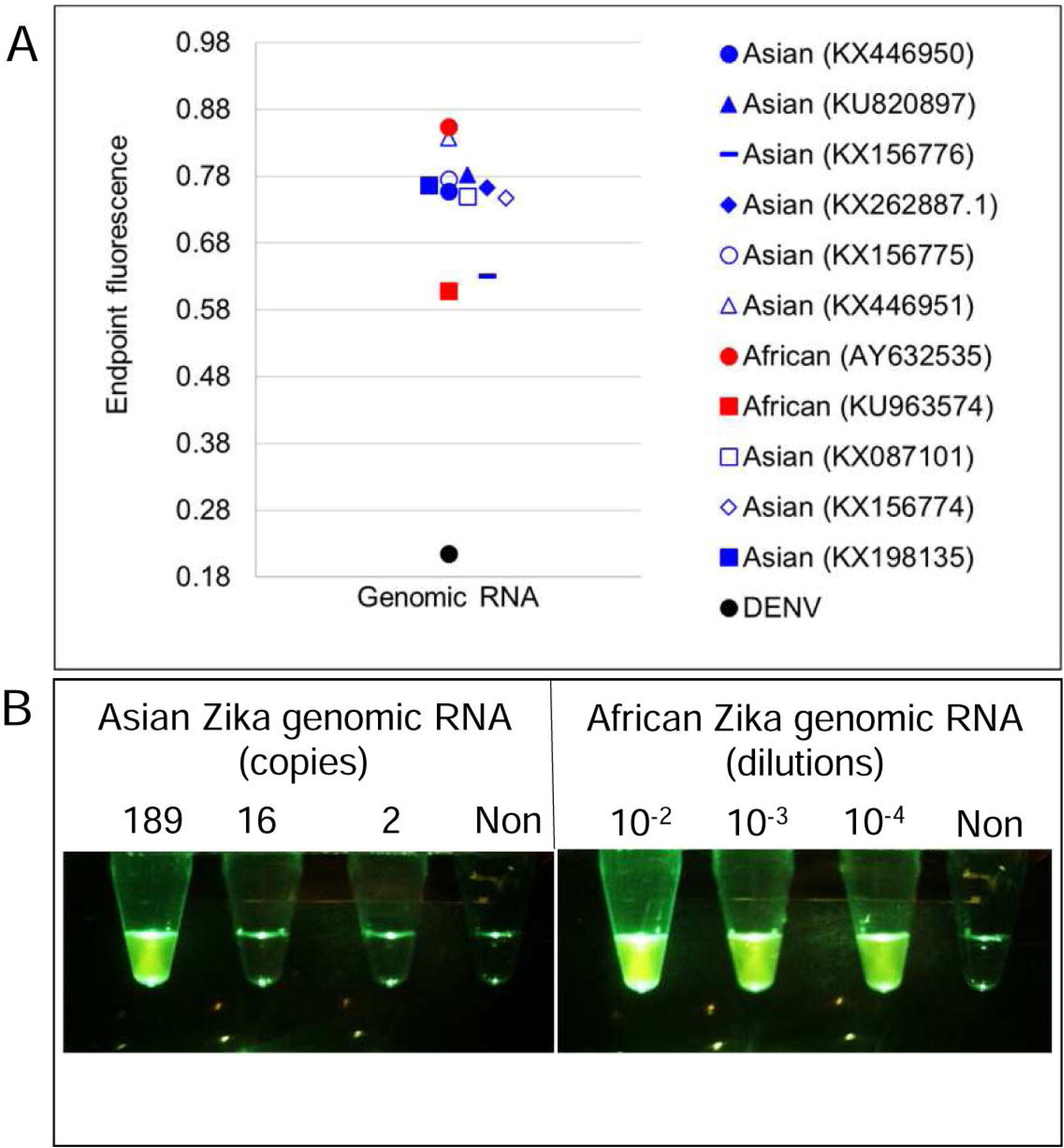
Detection of Asian and African lineage ZIKV genomes using degenerate 4-Plex-LAMP-2GO assays. (A) Whole cell RNA from cells infected with DENV, or Asian or African lineage ZIKV (indicated by their GenBank accession numbers) were used as templates for amplification. 2GO probe fluorescence signals measured at amplification endpoint using LightCycler 96 real-time PCR machine are depicted as blue (Asian ZIKV), red (African ZIKV), and black (DENV) markers. (B) Detection limit of degenerate 4-Plex-LAMP-2GO assay for ZIKV genomic RNA. Indicated copies of an Asian lineage ZIKV genome (left panel), indicated dilutions of an African ZIKV genome (right panel), and non-specific DENV genomes (‘Non’) were amplified using 4-Plex-LAMP-2GO assays. 2GO probe fluorescence was imaged at amplification endpoint using a cellphone. For all experiments, representative results from three replicate tests are depicted.

### Identification of ZIKV-infected mosquitoes using cellphones and smart molecular diagnostics

To evaluate assay performance in natural biological matrices, *Ae. aegypti* mosquitoes were fed a blood meal spiked with ZIKV virions and then individually tested for viral nucleic acids after 14 days. Half of these mosquitoes that were exposed to ZIKV generated bright endpoint fluorescence in both singleplex (NS1 LAMP-OSD and Capsid LAMP-OSD) and multiplex (4-plex-LAMP-2GO) degenerate reverse transcription LAMP-based assays (**Figure 8, Supplementary Figure 2**). Parallel tests performed in the absence of primers, and hence absence of LAMP amplification and concomitant probe activation, revealed negligible signal inflation due to sample auto-fluorescence, indicating the positive LAMP signal was due to these mosquitoes developing a viral infection. These mosquitoes also tested positive by ZIKV NS2b TaqMan qRT-PCR assay and were found to contain ~6×10^3^ to 4×10^5^ NS2b RNA copies per mosquito (**Figure 8**). Since only a 1/33 fraction of each mosquito was used for a LAMP test, these results suggest that the LAMP-based assays were able to generate a readily discernable bright visible fluorescence starting from as few as 190 copies of ZIKV genomic material embedded in crude virus-infected mosquito matrix.

**Figure 8.**
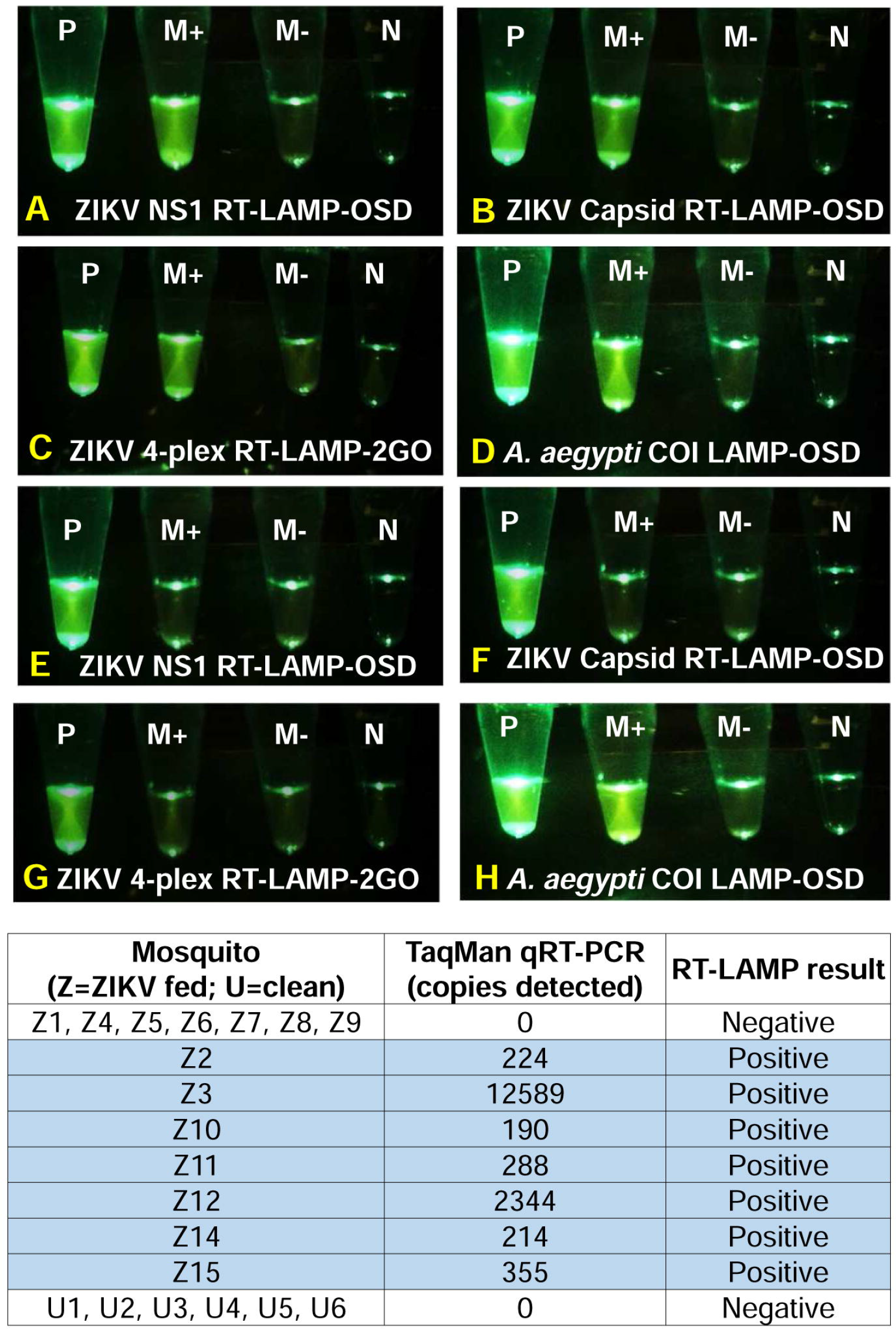
Detection of Zika virus-infected mosquitoes using single- and multiplex degenerate reverse transcription LAMP assays. Zika virus-infected (panels A-D) and uninfected (panels EH) *Aedes aegypti* mosquitoes were directly analyzed using singleplex NS1 and capsid LAMP-OSD assays or with 4-plex-LAMP-2GO assays. As a positive control, mosquitoes were tested using the *A. aegypti coi* LAMP-OSD assay (panels D and H). Smartphone images acquired after 2h of amplification are depicted. P: positive control; M+: mosquito analyte with LAMP primers; M-: mosquito analyte without LAMP primers; N: no template control. Results of NS2b TaqMan qRT-PCR analysis of all mosquitoes are tabulated.

Capsid and NS1 LAMP-OSD assays and 4-Plex-LAMP-2GO assays of the remaining half of the ZIKV-fed mosquitoes remained as dark as primer-less assays (**Supplementary Figure 2**). Our LAMP-based assays were in 100% agreement with TaqMan qRT-PCR, which also failed to detect ZIKV nucleic acids in these ZIKV-fed mosquitoes (**Figure 8**). These results suggest an absence of detectable infection in these mosquitoes, which is consistent with previously reported infection rates.^34, 35, 36, 37, 38^ Specificity of LAMP-based assays was confirmed by analyzing 6 additional age-matched *Ae. aegypti* mosquitoes that had not received a ZIKV-spiked blood meal and were negative for ZIKV when tested by qRT-PCR. None of the uninfected mosquitoes triggered a false positive signal; LAMP assays containing these uninfected mosquitoes remained indistinguishable from primer-less assays (**Supplementary Figure 2**).

Absence of signal in ZIKV-uninfected mosquitoes was not due to inhibition of amplification. Both ZIKV-infected and uninfected mosquitoes generated bright fluorescence when tested using a LAMP-OSD assay for *Ae. aegypti* mitochondrial cytochrome oxidase I (*coi*) gene^39^ (**Figure 8, Supplementary Figure 2**).

These results indicate that our smart molecular diagnostic assays perform at par with qRT-PCR for direct analysis of ZIKV virions in infected mosquitoes. Similar to singleplex Capsid and NS1 LAMP-OSD assays, the multiplex 4-Plex-LAMP-2GO assays operated without discernible inhibition or spurious actuation in this complex milieu and demonstrated 100% specificity and 100% sensitivity for viral detection.

### Detection of Zika virus in human saliva using cellphones and smart molecular diagnostics

Saliva is a convenient, non-invasive clinical sample that is increasingly being used for first-line rapid detection of pathogens including ZIKV.^40, 41^ To demonstrate the utility of our smart molecular diagnostic system in rapid salivary diagnostics, surrogate clinical samples were prepared by spiking human saliva with different amounts of Asian lineage ZIKV virions (**Supplementary Figure 1**) followed by direct analysis using singleplex degenerate NS1 LAMP-OSD and degenerate 4-Plex-LAMP-2GO assays. To verify diagnostic accuracy, saliva samples spiked with an unrelated RNA virus were also tested. To ascertain diagnostic speed, assay fluorescence was imaged using a cellphone at various intervals.

Our results demonstrate that both the singleplex NS1 LAMP-OSD assay and the multiplex 4-Plex-LAMP-2GO assay could detect as few as 2 x 10^4^/ml of ZIKV virions in human saliva within 90 min (**Figure 9**). Since only a 1/200 fraction of this saliva sample was used in the nucleic acid tests, it is evident that our LAMP-based assays were able to generate positive signal from only 100 ZIKV virions. Saliva containing higher concentration of ZIKV virions yielded faster results with both assays. A typically encountered physiological viral load (2 x 10^5^ virions / ml) generated bright fluorescence in the singleplex NS1 LAMP-OSD assay within only 30 min. The multiplex 4-Plex-LAMP-2GO assay required 60 min to generate discernible signal with the same sample. Neither assay generated non-specific fluorescence when challenged with saliva containing as many as 10^7^/ml of non-specific virions.

**Figure 9.**
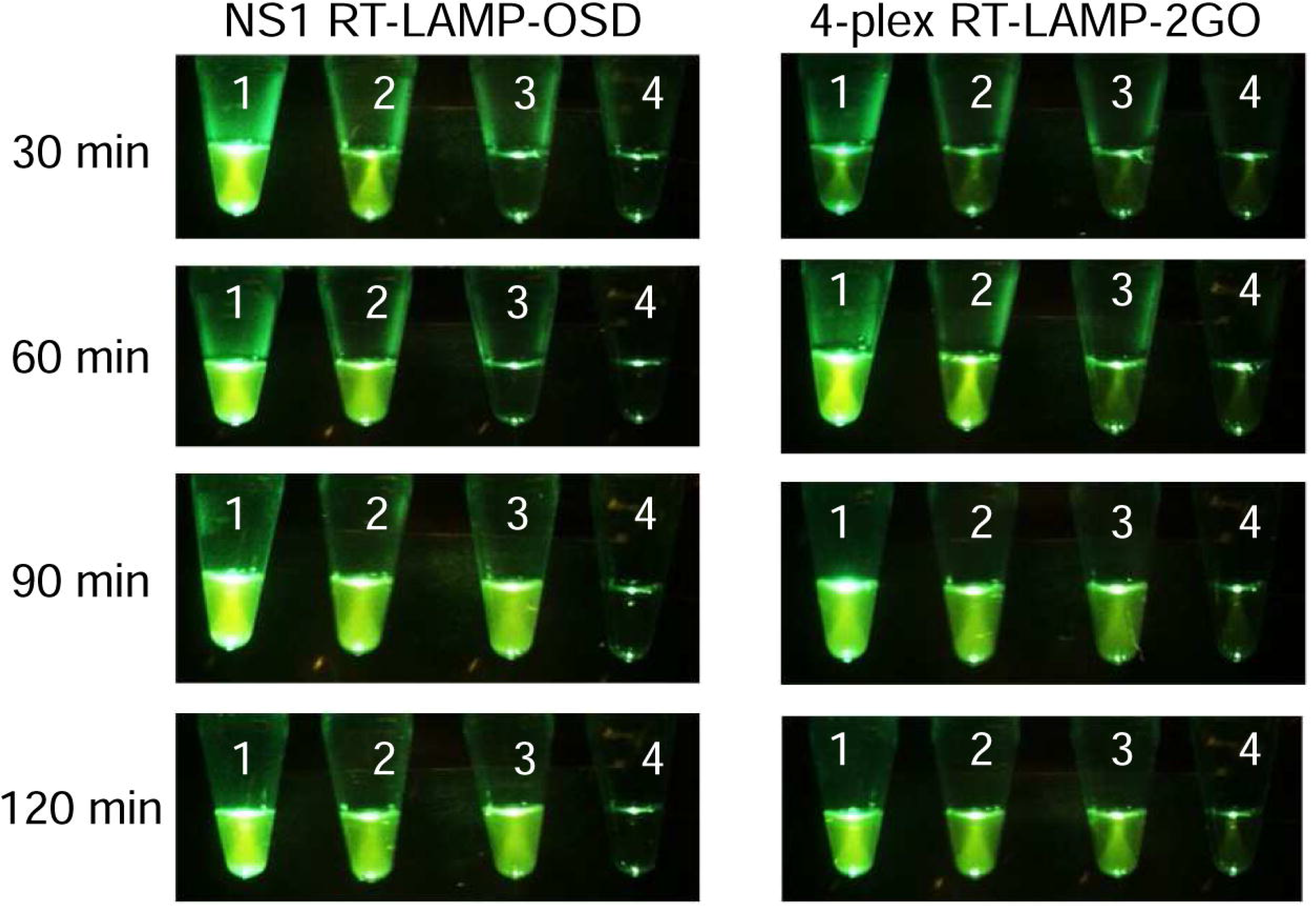
Detection of ZIKV virions in human saliva using single- and multiplex degenerate reverse transcription LAMP assays. Asian lineage ZIKV virions spiked in human saliva – 1: 2×10^6^ virions/ml; 2: 2×10^5^ virions/ml; 3: 2×10^4^ virions/ml; 4: 10^7^ non-specific virions/ml – were detected using direct singleplex analysis with degenerate NS1-specific LAMP-OSD assays or with multiplex degenerate 4-Plex-LAMP-2GO assays. Smartphone images of OSD or 2GO probe fluorescence acquired after different durations of amplification are depicted.

These results demonstrate the ability of our smart molecular diagnostic platform to rapidly amplify and signal the presence of target nucleic acids in human saliva without interference from spurious signal.

## Discussion

While iNAATs have potential as POC diagnostics, they lack the versatility and intelligence to deal with genetically variable and rapidly evolving targets. To fill this need we have created a smart molecular diagnostic composed of a parallel set of degenerate primers that can readily amplify variant targets and an integrated multiplex OR-gated strand exchange logic processor that ‘computes’ and returns the presence of target relative to even closely related non-target sequences. The fact that multiple, different amplification reactions can be carried out in parallel, and yet the user need only attend to the final, fluorescent signal highlights the power of the diagnostic, and presages the possibility of even more complex integration of molecular signals. For example, strand exchange nucleic acid computation has allowed (i) analysis of repetitive DNA targets of greater than 500 nucleotides in length with single nucleotide specificity, using ‘modular probes’ composed of multiple short oligonucleotides that undergo competitive hybridization;^42^ (ii) sensing and imaging of nucleic acids, enzymes, small molecules, proteins, and tumor cells via hairpin oligonucleotide probes and the hybridization chain reaction (HCR);^43^ and (iii) coupling nucleic acid detection to translational reporters via amplicon-mediated strand displacement reactions that couple to reporter protein production in a cell-free translation system.

More importantly, the robustness of our smart molecular diagnostic allows it to perform with little or no error in a variety of inclement environments, including with crude preparations of macerated mosquitoes. LAMP-based smart molecular assays of ZIKV-infected and un-infected mosquitoes could be performed with results that were on par with those obtained by qPCR analyses. As few as 190 copies of ZIKV RNA could be readily identified using 4-Plex-LAMP-2GO assays without any false positive reactions. Similarly, both single and multiplex LAMP-based assays could identify 2 x 10^4^/ml ZIKV virions spiked in human saliva within 90 min. These assays were equally adept at identifying both Asian and African lineages of ZIKV without cross-reaction with related or co-circulating viruses. The assay thus finds itself in a relatively rarified set of molecular diagnostics that are truly capable of performing in demanding field conditions. For instance, RPA has demonstrated near 100% sensitivity and specificity in field tests for detection of many viruses, bacteria, and parasites.^44, 45, 46^ Similarly, the isothermal nicking enzyme amplification reaction (NEAR) automated on the Clinical Laboratory Improvement Amendments (CLIA)-waived portable device Alere i (Alere, Scarborough, MA) is being used for POC diagnosis of infectious agents such as influenza viruses.^47^

Overall, though, our envisioned smart molecular diagnostics platform has much greater versatility for field use going forward. For instance, most field applications of RPA involve purification of nucleic acid analytes from clinical specimens. In contrast, our one-pot reactions can directly analyze crude biospecimens without necessitating complicated sample preparation and nucleic acid purification. Field applications of RPA and NEAR typically employ commercial instruments costing a few thousand dollars for assay readout^46^ whereas our assay requires a simple instrument-free visual ‘yes/no’ score for presence or absence of signal at endpoint.

Most importantly, existing iNAATs lack design considerations to counter target variability, a deficiency that may reduce sensitivity^47^ and in extreme cases lead to assay failure. For example, RPA uses a recombinase enzyme to facilitate invasion of double stranded templates by primers, which are then extended by a strand displacing DNA polymerase resulting in signal amplification.^6^ Amplicon detection is typically achieved using fluorophore-labeled probes that are measured using fluorimetry or converted to a color readout using lateral flow assays.^6, 48^ NEAR relies on DNA nicking and extension reactions using nicking enzymes and strand displacing DNA polymerase to achieve exponential amplification of target DNA or RNA.^49^ The resulting amplicons are typically converted to a fluorescence signal using cleavage probes. Neither configuration would be particularly suitable for degenerate priming and / or strand exchange computations, as opposed to our internally redundant Boolean logic processing assay.

In conclusion, our smart molecular diagnostic overcomes the principal challenges faced by most iNAATs – non-specific amplification, false negatives, and false positives – while maintaining an easy-to-use user-interface. This assay platform should be of greatest utility for the development of disposable point-of-care devices. Despite performing multi-component analytical operations, the molecular computations are hidden from the user in a one-pot reaction that can both directly receive the sample and emanate visual results. We have previously demonstrated that the ready-to-use LAMP-OSD assay mixes can be lyophilized and stored for extended durations without loss of activity.^13^ These various features should minimize device intricacies, and in turn reduce cost, especially since only two protein enzymes and a single fluorophore-quencher pair are required. Most importantly, the molecular probes can be configured to capture both current and future fast-evolving targets, such as the Zika viruses, with both greater breadth and greater accuracy than existing tests.

## Methods

### Chemicals and reagents

All chemicals were of analytical grade and were purchased from Sigma-Aldrich (St. Louis, MO, U.S.A.) unless otherwise indicated. All enzymes and related buffers were purchased from New England Biolabs (NEB, Ipswich, MA, USA) unless otherwise indicated. All oligonucleotides and gene blocks (summarized in **Supplementary Tables T1, T2, T3, and T5)** were obtained from Integrated DNA Technologies (IDT, Coralville, IA, USA). Virus genomic RNA and inactivated virions (**Supplementary Table T4**) were obtained from BEI Resources (Manassas, VA, USA).

### Cloning of gBlocks and PCR amplification of transcription templates

The gBlock double stranded DNA surrogates of ZIKV sequences were designed to include a T7 RNA polymerase promoter at their 5’-ends. These gBlocks were cloned into the pCR2.1-TOPO vector (Fisher Scientific, Hampton, NH) by Gibson assembly (NEB) according to the manufacturer’s instructions.^50^ Cloned plasmids were selected and maintained in an *E. coli* Top10 strain. Plasmid minipreps were prepared from these strains using Qiagen miniprep kit (Qiagen, Valencia, CA, USA). All gBlock inserts were verified by Sanger sequencing at the Institute of Cellular and Molecular Biology Core DNA Sequencing Facility.

For performing *in vitro* run-off transcription, ZIKV gene segments cloned in a pCR2.1-TOPO vector were amplified from sequenced plasmids by PCR using Phusion DNA polymerase. PCR products were verified by agarose gel electrophoresis and then purified using the Wizard SV gel and PCR Clean-up system (Promega, Madison, WI, USA), according to the manufacturer’s instructions.

### *In vitro* transcription

Some 1000 ng of purified linear, double-stranded DNA transcription templates were transcribed using the HiScribe T7 High Yield RNA synthesis kit (NEB) according to the manufacturer’s instructions. Transcription was allowed to occur at 37 °C for 2 h. Subsequently the transcription reactions were incubated with 2 units of DNase I (NEB) at 37 °C for 30 min to degrade the template DNA prior to RNA gel purification.

### Denaturing polyacrylamide gel electrophoresis and RNA gel purification

Denaturing 8% polyacrylamide gels containing 7 M urea were prepared using 40% acrylamide and bis-acrylamide solution, 19:1 (Bio-Rad) in 1X TBE buffer (89 mM Tris Base, 89 mM Boric acid, 2 mM EDTA, pH 8.0) containing 0.04% ammonium persulphate and 0.1% TEMED. An equal volume of 2X denaturing dye (7 M urea, 1X TBE, 0.1% bromophenol blue) was added to the RNA samples. These were incubated at 65 °C for 3 min followed by cooling to room temperature before electrophoresis. RNA bands were demarcated using UV shadowing. Desired bands were excised from the gel and the RNA was eluted twice into TE (10:1, pH 7.5) buffer (10 mM Tris-HCl, pH 7.5, 1 mM EDTA, pH 8.0) by incubation at 70 °C and 1000 rpm for 20 min. Acrylamide traces were removed by filtering eluates through Ultrafree-MC centrifugal filter units (EMD Millipore, Billerica, MA, USA) followed by precipitation with 2X volume of 100% ethanol in the presence of both 15 µg GlycoBlue (Life Technologies) and 0.3 M sodium acetate, pH 5.2. RNA pellets were washed once in 70% ethanol. Dried pellets of purified RNA were resuspended in 0.1 mM EDTA and stored at −80 °C.

### LAMP primer and strand exchange probe design

ZIKV genomic sequences were analyzed for variation using the National Center for Biotechnology Information (NCBI) Virus Variation Resource.^51^ ZIKV sequences were also compared to related and co-circulating viruses, such as DENV and CHIKV, using MUSCLE.^52, 53^ Four relatively conserved genomic regions in ZIKV capsid, NS1, NS3, and NS5 genes were chosen for primer design. The Primer Explorer v5 LAMP primer design software (Eiken Chemical Co., Japan) was used for generating LAMP primer sets composed of the outer primers F3 and B3 and the inner primers FIP and BIP. Primer design was constrained to include at least a 40 base pair (bp) gap between the F1 and F2 as well as between the B1 and B2 priming sites. Loop primers and stem primers were manually designed. Primer specificity for ZIKV isolates and a corresponding lack of significant cross-reactivity to other nucleic acids of human or pathogenic origin was further assessed using NCBI BLAST.^54, 55^ Polymorphic loci in the primers were substituted with degenerate bases although, some polymorphisms near the 5’-end of F3 or B3 or near the middle of FIP or BIP were ignored to reduce the degeneracy burden and to ensure efficient amplification.^15^ Our previous work had shown that such mismatches minimally disrupted LAMP primer efficiency.^15^

The nucleic acid circuit design software NUPACK^56^ was used to design four OSD probes, each specific to one of the four ZIKV amplicons, according to our previously published design rules.^12^ Unique, fluorophore-labeled OSD strands were designed to bind between B1 and B2 sequences of NS5 and NS3 amplicons and between F1 and F2 sequences of NS1 and capsid amplicons. The quencher-labeled OSD strands were designed to be partially complementary to the fluorophore-labeled strand. Single-stranded toeholds at the 3’-end or 5’-end of fluorophore-labeled strands were designed to be 10 or 13 nucleotides long. All 3’-OH ends were blocked with inverted deoxythymidine (dT) to prevent extension by DNA polymerase. Polymorphic loci in all four OSD probes were substituted with appropriate degenerate bases. These degenerate OSD probe strands were then used as a basis for designing the two-input 2GO and four-input 4GO probes. NUPACK was used to visualize and optimize probe architecture with variables such as buffer composition, temperature, and oligonucleotide concentrations.

### Assembly of strand exchange probes

OSD probes were prepared by annealing 1 µM of the fluorophore-labeled OSD strand with 5 µM of the quencher-labeled strand in 1X Isothermal buffer (NEB: 20 mM Tris-HCl, 10 mM (NH_4_)_2_SO_4_, 50 mM KCl, 2 mM MgSO_4_, 0.1% Tween 20, pH 8.8 at 25°C). Annealing was performed by denaturing the oligonucleotide mix at 95 °C for 1 min followed by slow cooling at the rate of 0.1 °C/s to 25 °C. Excess annealed probe was stored at −20 °C.

The CAN3.2GO probe was assembled by annealing 4 µM of CAN3.2GO.Gate oligonucleotide with 6 µM of the quencher-labeled CAN3.2GO.Q oligonucleotide and 2 µM of the fluorophore-labeled CAN3.2GO.FAM oligonucleotide. Similarly, the N1N5.2GO probe was assembled by annealing 6 µM of N5N1.2GO.Gate oligonucleotide with 8 µM of the quencher-labeled N5N1.2GO.Q oligonucleotide and 2 µM of the fluorophore-labeled N5N1.2GO.FAM oligonucleotide. The three oligonucleotide components of each 2GO probe were mixed in 1X Isothermal buffer supplemented with 8 mM MgSO_4_ and incubated for 1 min at 95 ºC. The mixture was then slowly cooled at the rate of 0.1 °C/s to 25 °C. Excess annealed 2GO probes were stored at −20 °C.

The 4GO probe was assembled by annealing 4 µM of the 4GO.S1 strand, 1 µM of the fluorophore-labeled 4GO.S2 strand, 4 µM of the quencher-labeled 4GO.S3 strand, 4 µM of the 4GO.S4 strand, and 3 µM of the 4GO.S5 strand. All oligonucleotides were mixed in 1X Isothermal buffer supplemented with 8 mM MgSO_4_ and incubated for 1 min at 95 °C. The mixture was then slowly cooled at the rate of 0.1 °C/s to 25 °C. Excess annealed 4GO probes were stored at −20 °C.

### One-pot RT-LAMP assay

All one-pot RT-LAMP assays were assembled in 25 µL reactions containing 1X Isothermal buffer supplemented with 1.4 mM deoxyribonucleotides (dNTPs), 0.4 M betaine, 6 mM additional MgSO_4_, 16 units of Bst 2.0 DNA polymerase (NEB), and 7.5 units of warmstart RTx reverse transcriptase (NEB). Primer-containing singleplex reactions were appended with 2.4 µM each of a single type of FIP and BIP primer, 1.2 µM of the corresponding loop primer, 1.2 µM of the corresponding stem primer (only for the capsid assay), and 0.6 µM each of the corresponding F3 and B3 primer. Primer-containing 4-Plex LAMP reactions received 0.6 µM each of the four FIP and BIP primers, 0.3 µM of each of the four loop primers, 0.3 µM of the capsid stem primer, and 0.15 µM each of the four F3 and B3 primers. The total primer content of these multiplex assays was the same as individual primer concentrations used in singleplex reactions. Control assays lacking primers received equivalent volume of TE 10:0.1 buffer (10 mM Tris, pH 7.5 and 0.1 mM EDTA pH 8.0).

Singleplex assays read using OSD reporters received 200 nM of the fluorophore-labeled strand annealed with a five-fold excess of the complementary quencher-labeled strand (see subsection “*Assembly of strand exchange probes”*). 4-Plex LAMP-OSD assays received 50 nM of each of the four fluorophore-labeled OSD strands pre-annealed individually with five-fold excess of their corresponding quencher-labeled complementary strands.

Assays analyzed using 2GO probes were supplemented with 50 mM trehalose and 100 nM of the fluorophore-labeled probe strand pre-annealed with the complementary gate and quencher-labeled oligonucleotides as described in sub-section “*Assembly of strand exchange probes”*. 4-Plex assays read using the complete bipartite signal transducer contained 100 nM each of both CAN3.2GO and N1N5.2GO probes. Assays read using 4GO probes were also supplemented with 50 mM trehalose along with 80 nM of the fluorophore-labeled strand annealed with the remaining four strands of the 4GO probe as described in the sub-section “*Assembly of strand exchange probes”*.

Real-time measurement of fluorescence accumulation was performed using the LightCycler96 real-time PCR machine (Roche, Basel, Switzerland). Reactions were subjected to isothermal amplification by programming 45 cycles of 150 sec at 65 °C (step 1) followed by 30 sec at 65 °C (step 2). Fluorescence was measured in the FAM channel during step 2 of each amplification cycle. Post-amplification reporter fluorescence was also monitored by programming the LightCycler96 to hold the assays at 37 °C for 40 min.

LAMP-based assays intended for visual readout and smartphone imaging were assembled in 0.2 ml optically clear thin-walled tubes with low auto-fluorescence (Axygen, Union City, CA, USA). Following 90 – 120 min of amplification at 65 °C, the reactions were imaged at room temperature using an unmodified iPhone 6 and an UltraSlim-LED transilluminator (Syngene, Frederick, MD, USA). In some experiments, our previously described in-house 3D-printed imaging device^13^ was used for fluorescence visualization and smartphone imaging. Briefly, this device uses Super Bright Blue 5 mm light emitting diodes (LED) (Adafruit, New York, NY, USA) to excite fluorescence. Two cut-to-fit layers of inexpensive >500 nm bandpass orange lighting gel sheets (Lee Filters, Burbank, CA, USA) on the observation window filter the fluorescence for observation and imaging.

The following ZIKV templates were analyzed by LAMP-OSD, LAMP-2GO, and LAMP-4GO assays: (i) zero to several thousands of copies of ZIKV synthetic RNA or genomic RNA in TE 10:0.1 buffer; (ii) ZIKV virions in TE 10:0.1 buffer and in human saliva (see sub-section “*Analysis of ZIKV virions*”); (iii) Un-infected and ZIKV-infected *Ae. aegypti* mosquitoes (see subsection “*Rearing and analysis of ZIKV-infected mosquitoes*”). Negative controls included (i) assays without any templates; (ii) assays with non-specific templates including synthetic RNA, DENV genomic RNA, and CHIKV genomic RNA; and (iii) assays with non-specific amplicons (generated by substituting with a non-specific template and its cognate LAMP primer set).

### TaqMan qRT-PCR analysis

The previously reported ZIKV NS2b-specific TaqMan qRT-PCR assay (**Supplementary Table T5**)^33^ was performed using either the Evoscript RNA Probes Master (Roche, Basel, Switzerland) or the TaqMan RNA-to-Ct 1-Step Kit (Thermo Scientific, Waltham MA, USA). Standard curves were prepared using a ten-fold dilution series of 10^5^ to 10 copies of a synthetic RNA template (**Supplementary Table T5**). Reactions without any templates served as negative controls. These templates were subjected to one-step qRT-PCR in 10 µL reactions containing 800 nM each of forward (Zika4481_F) and reverse (Zika4552c_R) primers along with 200 nM of the TaqMan probe (Zika4507cTqMFAM). Amplification was performed by adding either (i) 2 µL of the 5X Evoscript RNA probes Master or (ii) 0.25 µL of the 40X TaqMan RT Enzyme Mix (Thermo Scientific) along with 5 µL of the 2X TaqMan RNA-to-Ct 1-Step master mix. Reactions were incubated for 30 min at 60 °C followed by 10 min at 95 °C. Subsequently 45 cycles of 15 sec at 95 °C and 30 sec at 55 °C were performed. TaqMan fluorescence was measured during the second step of each cycle. Reactions containing Evoscript RNA Probes Master were analyzed using the LightCycler96 qPCR machine while TaqMan RNA-to-Ct 1-Step assays were performed on the StepOnePlus real-time PCR machine (Thermo Scientific).

### Analysis of ZIKV virions

Heat-inactivated virions of Asian ZIKV strain PRVABC59 (BEI Resources, Manassas, VA, USA) were used as templates for LAMP-based assays and for one-step qRT-PCR. Virions were either diluted in TE 10:0.1 buffer or in normal human saliva pooled from multiple donors (Catalog number: 991-05-P-5; Lee Biosolutions, Maryland Heights, MO, USA) prior to being directly added to LAMP-based and qRT-PCR reactions. The human saliva was heated to 95 °C for 15 min and then cooled prior to spiking with virions.

### Rearing and analysis of ZIKV-infected mosquitoes

*Ae. aegypti* mosquitoes were reared under conventional conditions in the insectary at the University of Texas medical Branch, Galveston, TX, USA. To obtain ZIKV-infected insects, mosquitoes were starved for a period of 24 hours then offered a sheep blood meal (Colorado Serum Company, Denver, CO, USA) containing 10^6^ virions using a hemotek membrane system (Hemotek). Un-infected mosquitoes received a blood meal that did not contain virus. After 14 days, mosquitoes were collected and immediately frozen at −80 °C. Prior to molecular testing, each mosquito was heated for 10 min at 60 °C in order to inactivate the Zika virus^57^ for compliance with biosafety level 2 requirements. Each mosquito was then manually crushed in a 1.7 mL microcentrifuge tube using a disposable micropestle (Fisherbrand™ RNase-Free Disposable Pellet Pestles, Catalog number 12-141-364, Fisher Scientific, Hampton, NH, USA). Each macerated mosquito was re-suspended in 100 µL water. A 2 µL aliquot of this mosquito sample was directly assessed by *Ae. aegypti coi* LAMP-OSD assay in both primer-containing and primer-free reactions. For ZIKV LAMP-based reactions and qRT-PCR assays, 3 µL mosquito samples were directly used as sources of templates. Both primer-containing and primer-less LAMP-based assays were operated.

## Acknowledgements

This work was supported by a Bill and Melinda Gates Foundation grant (OPP1128792) and the National Science Foundation BEACON (DBI-0939454) grant to ADE and a NIAID-SBIR (R43 AI131948) grant to ADE and GLH. GLH was also supported by the Wolfson Foundation and Royal Society, NIH grants (R21AI138074, R21AI124452 and R21AI129507), a University of Texas Rising Star award, the Western Gulf Center of Excellence for Vector-borne Diseases (CDC grant CK17-005), the Robert J. and Helen Kleberg Foundation and the Gulf Coast Consortia. MS was supported by an NIH T32 fellowship (2T32AI007526). The papers contents are solely the responsibility of the authors and do not necessarily represent the official views of the Centers for Disease Control and Prevention or the Department of Health and Human Services.

## Electronic supplementary material

Supplementary information

